# Evidence of Absence Treated as Absence of Evidence: The Effects of Variation in the Number and Distribution of Gaps Treated as Missing Data on the Results of Standard Maximum Likelihood Analysis

**DOI:** 10.1101/755009

**Authors:** Denis Jacob Machado, Santiago Castroviejo-Fisher, Taran Grant

**Affiliations:** University of North Carolina at Charlotte, College of Computing and Informatics, Department of Bioinformatcis and Genomics, 9201 University City Blvd, Charlotte–NC 28223, USA; Universidade de São Paulo, Programa Interunidades de Pós-Graduação em Bioinformática. Rua do Matão, 1010, CEP: 05508-090, São Paulo–SP, Brazil; Pontifícia Universidade Católica do Rio Grande do Sul, Laboratório de Sistemática de Vertebrados. Avenida Ipiranga, 6681, prédio 12, Partenon, CEP: 90619-900, Porto Alegre–RS, Brazil; Universidade de São Paulo, Instituto de Biociências, Departamento de Zoologia, Laboratório de Anfíbios. Rua do Matão, tv. 14, 101, Cidade Universitária, CEP: 05508-090, São Paulo–SP, Brazil

**Keywords:** alignment, ambiguous characters, indel, maximum likelihood, missing data, phylogenetic analysis

## Abstract

We evaluated the effects of variation in the number and distribution of gaps (i.e., no base; coded as IUPAC “.” or “–”) treated as missing data (i.e., any base, coded as “?” or IUPAC “N”) in standard maximum likelihood (ML) analysis. We obtained alignments with variable numbers and arrangements of gaps by aligning seven diverse empirical datasets under different gap opening costs using MAFFT. We selected the optimal substitution model for each alignment using the corrected Akaike Information Criterion (AICc) in jModelTest2 and searched for the optimal trees for each alignment using default search parameters and the selected models in GARLI. We also employed a Monte Carlo approach to randomly insert gaps (treated as missing data) into an empirical dataset to understand more precisely the effects of their variable numbers and distributions. To compare alignments quantitatively, we used several measures to quantify the number and distribution of gaps in all alignments (e.g., alignment length, total number of gaps, total number of characters containing gaps, number of gap openings). We then used these variables to derive four indices (ranging from 0 to 1) that summarize the distribution of gaps both within and among terminals, including an index that takes into account their optimization on the tree. Our most important observation is that ML scores correlate negatively with gap opening costs, and the amount of missing data. These variables also cause unpredictable effects on tree topologies. We discuss the implications of our results for the traditional and tree-alignment approaches in ML.

## Introduction

Standard maximum likelihood (ML) analysis of DNA sequences follows a three-step procedure composed of (I) multiple sequence alignment (MSA) using programs such as CLUSTAL X (1), MAFFT (2, 3), or MUSCLE (4), (II) substitution model selection using programs like jModelTest (5) or PartitionFinder (6), and (III) tree searching using, for example, GARLI (7), PhyML (8), RAxML (9), or IQ-Tree (10). In the first step, insertion/deletion (indel) events are inferred according to user-specified indel opening and extension costs and nucleotides inferred to be absent due to indels are represented in the alignment as gaps (coded as IUPAC “–”). In the second and third steps, gaps are treated as missing nucleotides and coded as ambiguities in the matrix (nucleotides of unknown identity; “?” or IUPAC “N”), thereby recasting evidence of absence as absence of evidence.

The effects of increasing amounts of ambiguity due to missing data are reasonably well understood: ML scores increase, the likelihood surface flattens, and, depending on the number and distribution of the ambiguities, topological relationships can change and support values can become inflated (11–16). Similarly, previous studies have examined the degree to which different methods of alignment can alter results (e.g., 17–22). However, to date no study has systematically investigated the effects of variation in the number and distribution of gaps treated as missing data on the results of standard ML analyses. Here, we evaluated the effects of this variation on model selection, ML score, and topology by analyzing highly variable alignments obtained by aligning diverse empirical datasets under different gap opening costs and by randomly replacing nucleotides with gaps.

## Methods

### Datasets, Model Selection, and Phylogenetic Analysis

We summarized the analyses of the eight empirical datasets in Fig. 1. Analyses varied extensively in the number of terminals and sequence length (Table 1). We aligned datasets 1–7 using the program MAFFT v7.147b. We performed the alignment of these seven datasets using the Needleman-Wunsch algorithm and 1,000 cycles of iterative refinement (using the arguments “–globalpair” and “–maxiterate 1000”). We set the gap opening penalty to 0 and also to all values in a geometric progression of ratio 2 from 0.095625 to 24.48, including the program default value of 1.53. This resulted in 10 alignments per dataset that varied greatly in the number and distribution of gaps. Next, we converted the resulting MAFFT alignments from FASTA to NEXUS format using Mesquite v2.75 (build 564; 23) and selected the optimal substitution model for each alignment using the corrected Akaike Information Criterion (AICc) in jModelTest v2.1.4. We then performed ML tree searches in GARLI v2.01 using default search parameters and assuming the optimal substitution model selected previously by jModelTest. Each tree search comprised 640 replicates spawned in parallel using a homemade Python script (sudoParallelGarli.py).

**Table 1.** Basic information of the eight datasets used in this study. * Effects of variation in the number and distribution of gaps treated as missing data; ** Simulations.

| Dataset | Reference | Marker | Length (bp) | # terminals | Analyses |
| --- | --- | --- | --- | --- | --- |
| 1 | Wheeler and Hayashi 1998 | 18S rRNA | 940–2020 | 32 | * |
| 2 | Wheeler and Hayashi 1998 | 28S rRNA | 336–652 | 28 | * |
| 3 | Healy et al. 2009 | 18S rRNA | 554–2189 | 58 | * |
| 4 | Healy et al. 2009 | 28S rRNA | 668–4279 | 58 | * |
| 5 | Wei et al. 2014 | 12S mtDNA | 437–1011 | 31 | * |
| 6 | Mauro et al. 2014 | 16S mtDNA | 1534–1635 | 55 | * |
| 7 | Pozzi et al. 2014 | mtDNA Control Region (CR) | 515–2295 | 77 | * |
| 8 | Blotto et al. 2013 | CytB | 338–1003 (385) | 88 (85) | ** |

**Fig. 1.**
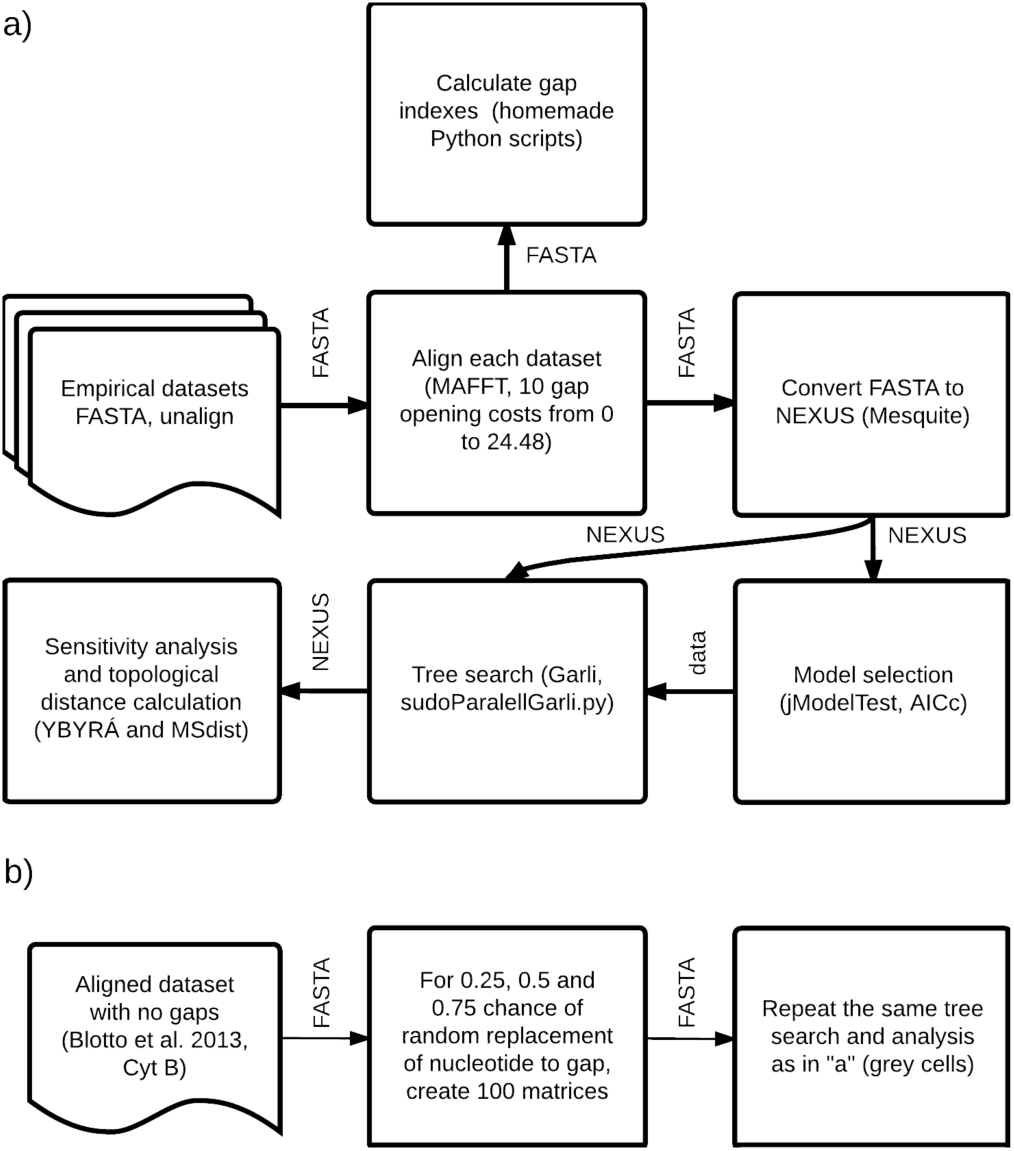
Summary of the methods for testing the effects of gaps treated as missing data in standard ML analysis. Datasets 8 is omitted since it was used for simulations with fixed alignment length. a) Empirical analysis with seven datasets (see Table 1). Analysis of 3,000 simulated matrices. Words by the arrows indicate the type of file or information being transferred. See text for additional details.

To clarify the effects of increasing amounts of ambiguity due to gaps and other possible alignment effects caused by the insertion of gaps during alignment, we performed additional Monte Carlo analyses using dataset 8 (Table 1). First, we aligned the data in MAFFT using the default gap opening cost of 1.53, trimmed the resulting alignment to include no leading or trailing gaps, and analyzed the matrix as described above. Next, we performed three rounds of 1,000 Monte Carlo replicates that randomly replaced nucleotides with gaps with probabilities of 0.25, 0.5, and 0.75, respectively, thereby increasing the number of gaps without altering nucleotide homology relationships or the length of the alignment (see below). Finally, we analyzed each of the resulting matrices in GARLI as described above, using the same substitution model selected for the original alignment.

All unaligned datasets and templates for the configuration files and execution scripts are available as supplementary material. We ran all compute-intensive analyses on the high performance computing cluster ACE, which is composed of 12 quad-socket AMD Opteron 6376 16-core 2.3-GHz CPU, 16MB cache, 6.4 GT/s compute nodes (= 768 cores total), eight with 128 GB RAM DDR3 1600 MHz (16 x 8GB), two with 256 GB (16 x 16GB), and two with 512 GB (32 x 16GB), and QDR 4x InfiniBand (32 GB/s) networking, housed at the Museu de Zoologia da Universidade de São Paulo (MZUSP).

### Alignment Characterization

We compared the 3,070 alignments generated for the eight datasets applying both the gap opening cost used to generate the alignments and several new measures and indices that describe the number and distribution of gaps in each alignment. For each alignment, we calculated alignment length, total number of gaps, total number of characters (columns, positions, transformation series) containing gaps, mean number of gap openings, and number of shared characteres (i.e., the number of columns that provide identical phylogenetic information.

Several indices and algorithms have been proposed to describe and compare multiple sequence alignments (e.g., 21, 24–27), some of which are derived from some of the same measures listed above. However, these methods focus on measuring overall genetic distances, structural modifications, or alignment accuracy or reliability, whereas we are specifically interested in evaluating the effects of varying the number and distribution of gaps. Consequently, we derived the following original indices (all defined to vary between 0–1) to summarize the distribution of gaps both within and among terminals.

#### Gap contiguity index (GCI)

GCI quantifies the degree to which gaps are grouped into contiguous strings or broken into short strings. For a given terminal with *g* gaps and *g*′ trailing gaps,

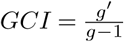

GCI is undefined if the sequence has no gaps or the sequence contains only a single trailing gap; otherwise, *GCI* = 0 if no gaps are contiguous and *GCI* = 1 if there is > 1 trailing gap and all gaps are contiguous. For an alignment-wide value, we report the mean GCI for all terminals in the alignment. Sequences without gaps are ignored during GCI calculations.

#### Nucleotide contiguity index (NCI)

NCI is equivalent to GCI but measures the contiguity of nucleotides instead of gaps. For a given terminal with *n* nucleotides and *n*′ trailing nucleotides,

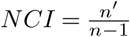

NCI is undefined if the alignment is composed of < 2 nucleotides; otherwise, *NCI* = 0 if all nucleotides for the terminal are separated by gaps and *NCI* = 1 if all nucleotides for the terminal are contiguous. For an alignment-wide value, we report the mean NCI for all terminals in the alignment.

#### Shared gaps index (SGI)

SGI quantifies the degree to which a given gap is shared among terminals. For a given character scored for *t* terminals of which *t*′ terminals possess a gap,

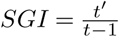

This index is undefined if the matrix contains only a single terminal; otherwise, *SGI* = 0 if the character contains no gaps and *SGI* = 1 if the gap is shared by all but one of the terminals (i.e., only one terminal possesses a nucleotide). For an alignment-wide value, we report the mean SGI for all characters that contain gaps (CG).

#### Topological gap index (TGI)

TGI summarizes the degree to which gaps are shared among terminals in the matrix. However, it ignores the topological distribution of those terminals – and, therefore, the topological distribution of the gaps – on the optimal tree. For a given character with gaps shared by *t*′ terminals and explained on the given tree by a minimum of *t*′′ gap↔nucleotide transformations, TGI is defined as

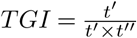

TGI is undefined if the character does not contain gaps; otherwise, *TGI* = 1 if a minimum of one gap↔nucleotide transformation explains the gaps in all terminals (i.e., a single split divides all the terminals that possess the gap from all the terminals that possess a nucleotide) and decreases as the minimum number of gap↔nucleotide transformations increases. For an alignment-wide value, we report the mean TGI for all characters that contain gaps.

### Evaluation Criteria

We evaluated the effects of variation in the number and distribution of gaps treated as missing data in standard ML phylogenetic analysis by comparing the alignment parameters (gap opening costs) and measures and indices with three response variables: (I) optimal substitution model selected using jModelTest; (II) the optimal ML score from GARLI; and (III) the optimal tree topology. To assess the effect on tree topology, we calculated the match split distances (MSD) between the optimal topologies using MS-dist v0.5 (28) and visualized their congruence using YBYRÁ (29). We used R v3.3.1 (30) to fit linear models for correlation analysis.

## Results

### Alignments

The different gap opening costs we used to align each dataset generated highly diverse alignments, as indicated by the variation in the values taken by all of the indices (Table S1). Among the alignments of sequences from datasets 1–7, mean GCI, mean NCI, mean SGI, and mean TGI values varied from 0.492–0.993, 0.375–0.992, 0.116– 0.758, and 0.394–0.855, respectively. The most variable datasets for each of our indices were dataset 5 for mean SGI (0.392–0.575), dataset 6 for mean GCI (0.492–0.935), and dataset 7 for mean NCI (0.790–0.992) and mean TGI (0.394– 0.571).

The 3,000 alignments generated by randomly replacing nucleotides with gaps in dataset 8 were also highly diverse. Alignment matrices composed of approximately 25% gaps had mean GCI, mean NCI, and mean SGI values of 0.234– 0.266, 0.741–0.761, and 0.245–0.260, respectively. Alignment matrices with approximately 50% gaps had mean GCI, mean NCI, and mean SGI values of 0.488–0.514, 0.488– 0.513, and 0.496–0.515, respectively. Lastly, alignment matrices with approximately 75% gaps had mean GCI, mean NCI, and mean SGI values of 0.738–0.758, 0.231–0.264, and 0.749–0.769, respectively.

### Model Selection

Despite the extensive variation among alignments, model selection varied little (Table 2). All models included gamma rate variation. Model selection chose the most complex model (GTR+I+G) for 80% of the alignments, including 100% of the alignments for datasets 3, 4, 6, and 7. Among the remaining datasets, we did not detect any trends in model selection. For example, dataset 5 varied most extensively, shifting between three models as gap opening costs increased: GTR+G for the two lowest gap opening costs, then HKY+G for the next three gap opening costs, GTR+G again for the next two, then GTR+I+G, GTR+G, and GTR+I+G for the three highest gap opening costs, respectively. In contrast, for dataset 2 the most complex model was chosen for the lowest two gap opening costs, then the less complex GTR+G, returning to the most complex model, then the even less complex HKY+G followed by GTR+G for the three highest gap opening costs.

**Table 2.** Models of nucleotide substitution for the 10 alignments of 7 datasets as selected by the AICc in jModelTest2. GTR = general time reversible mode; HKY = Hasegawa-Kishino-Yano model; SYM = symmetrical model; I = proportion of invariable sites; G = gamma distribution. * Alignments were performed in MAFFT. ** Default value.

| Gap opening cost* | Dataset 1 | Dataset 2 | Dataset 3 | Dataset 4 | Dataset 5 | Dataset 6 | Dataset 7 |
| --- | --- | --- | --- | --- | --- | --- | --- |
| 0 | GTR+I+G | GTR+I+G | GTR+I+G | GTR+I+G | GTR+G | GTR+I+G | GTR+I+G |
| 0.096 | GTR+I+G | GTR+I+G | GTR+I+G | GTR+I+G | GTR+G | GTR+I+G | GTR+I+G |
| 0.191 | GTR+I+G | GTR+G | GTR+I+G | GTR+I+G | HKY+G | GTR+I+G | GTR+I+G |
| 0.383 | GTR+I+G | GTR+I+G | GTR+I+G | GTR+I+G | HKY+G | GTR+I+G | GTR+I+G |
| 0.765 | GTR+I+G | GTR+I+G | GTR+I+G | GTR+I+G | HKY+G | GTR+I+G | GTR+I+G |
| 1.53** | GTR+I+G | GTR+I+G | GTR+I+G | GTR+I+G | GTR+G | GTR+I+G | GTR+I+G |
| 3.06 | SYM+I+G | HKY+G | GTR+I+G | GTR+I+G | GTR+G | GTR+I+G | GTR+I+G |
| 6.12 | GTR+I+G | GTR+G | GTR+I+G | GTR+I+G | GTR+I+G | GTR+I+G | GTR+I+G |
| 12.24 | GTR+I+G | GTR+G | GTR+I+G | GTR+I+G | GTR+G | GTR+I+G | GTR+I+G |
| 24.48 | GTR+I+G | GTR+G | GTR+I+G | GTR+I+G | GTR+I+G | GTR+I+G | GTR+I+G |

### Tree Topology

Although variation in the number and distribution of gaps treated as missing data had little effect on model selection, it had a substantial effect on tree topology (Fig. 2). Nevertheless, we did not detect any pattern to explain the observed variation in tree topology, and in many cases the most distant topologies were derived from alignments with adjacent gap opening scores, i.e., significant differences in tree topology were obtained with only minor variations in the alignment parameters and resulting quantity and distribution of gaps, even when there was no variation in the substitution model.

**Fig. 2.**
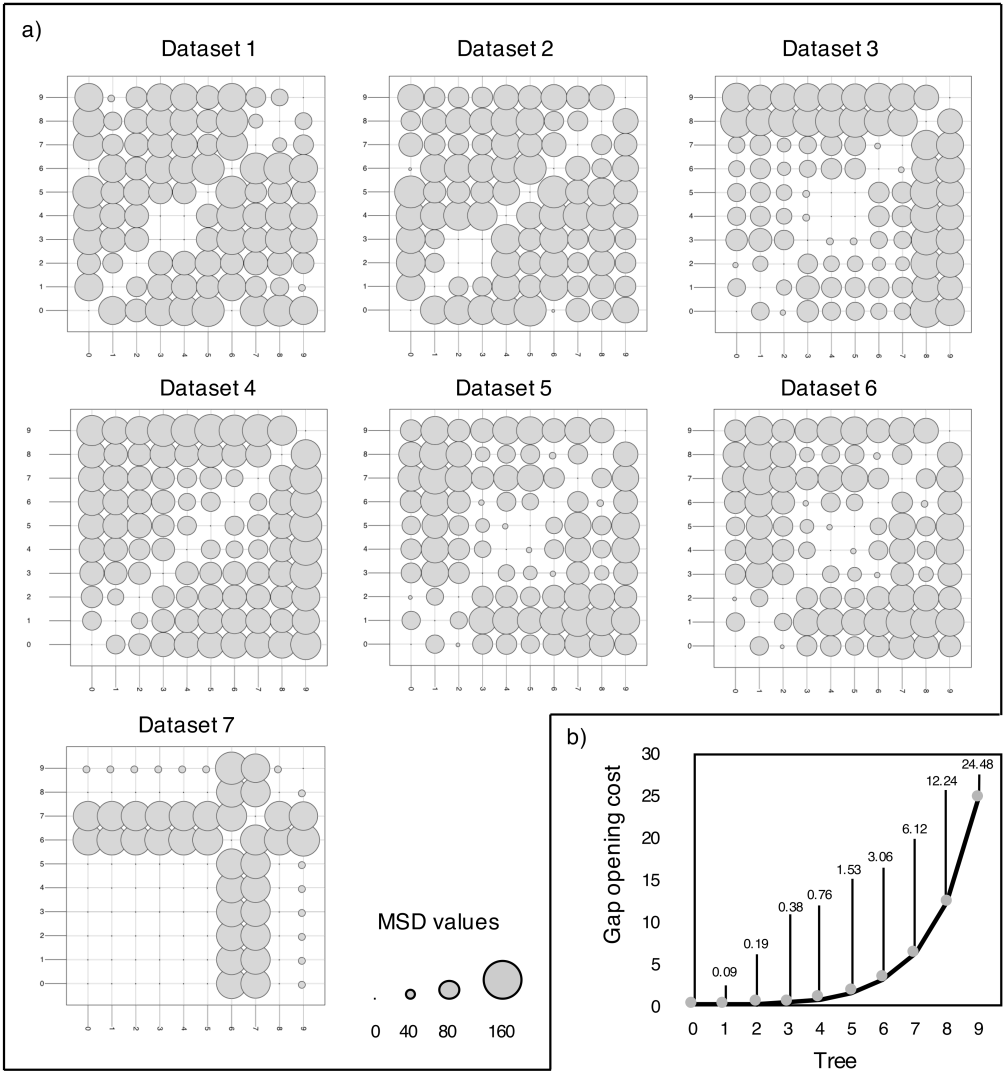
Variation in tree topology according to sequence alignment in datasets with variable alignment length (1 to 7). a) Visualization of the topological distance between each pair of trees, organized in ascending order of the gap opening cost used for multiple sequence alignment (0 to 9). The larger the circle, the higher the average match-split distance (MSD); b) Gap opening costs used during multiple sequence alignment (0 to 24.48) and the corresponding tree number (0 to 9).

### ML Score

For most datasets, the ML score negatively correlated with gap opening cost (adjusted *R*^2^ > 0.99; Table S2). For all datasets except dataset 4, ML score was also negatively correlated with GCI (adjusted *R*^2^ = 0.73–0.99) and NCI (adjusted *R*^2^ = 0.71–0.91), both of which measure the degree to which sequences form contiguous strings or are broken into short strings. In contrast, the ML score positively correlated with alignment length, percentage of gaps, and mean SGI in all datasets except dataset 4 (Fig. 3).

**Fig. 3.**
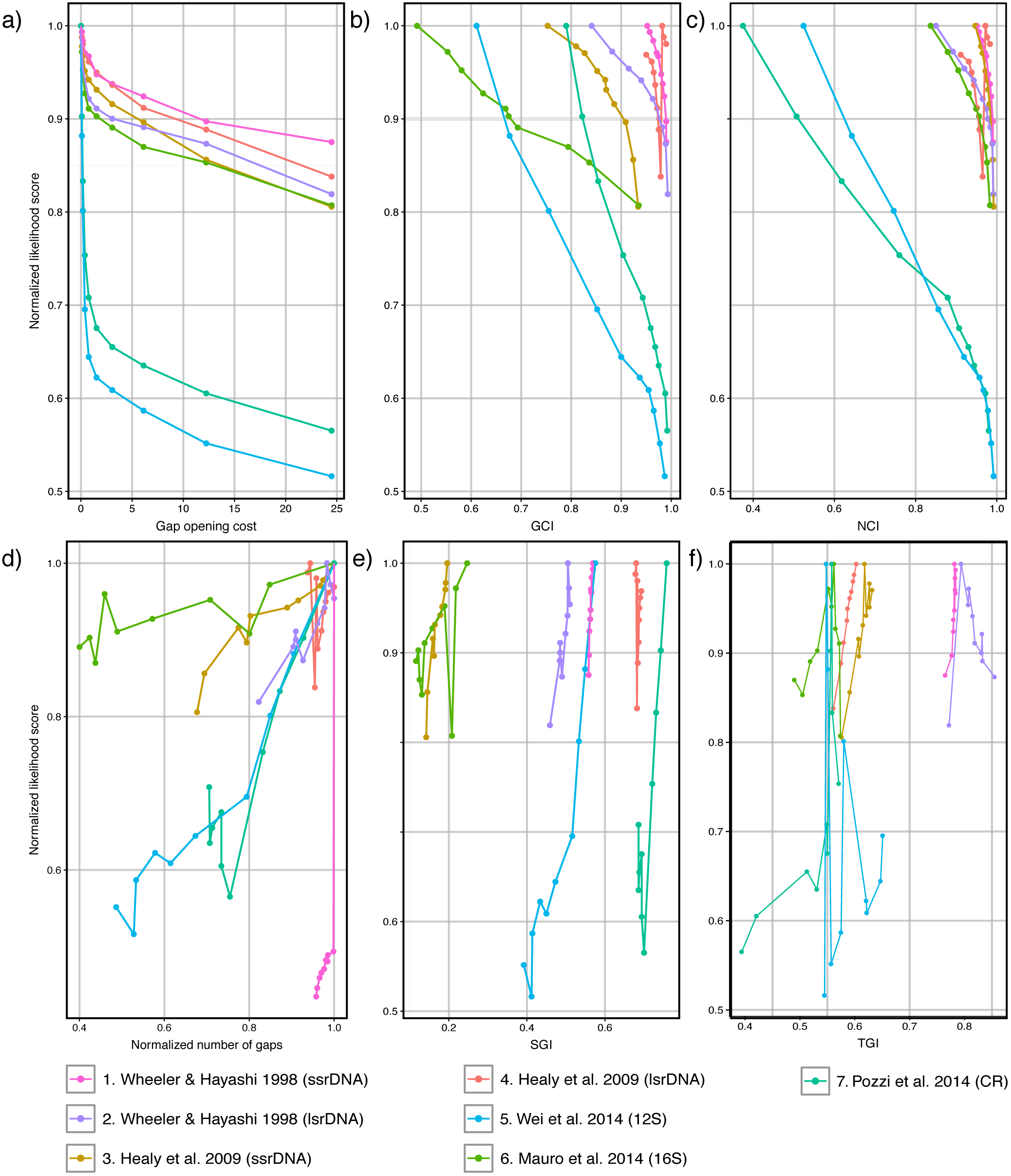
Variation of different variables as a result of changes in the gap opening score and the likelihood score of the corresponding tree. The Y-axis shows the normalized likelihood scores. The X-axis shows a) the gap opening cost, b) the average gap contiguity index (mean GCI), c) the average nucleotide contiguity index (mean NCI), d) the normalized number of gaps (percentage), e) the average shared gap index (mean SGI), and f) the average topological gap index (mean TGI). The variable length was omitted since it closely resembles the variable percentage of gaps. Analyzed data and the results of the linear model analyses are available at Tables S1 and S2, respectively.

Dataset 4 differed from all others in that the number of shared characters decreased as the gap opening cost increased (Fig. 4). The correlation analysis of the ML score and the mean TGI of dataset 4 had *R*^2^ = 0.98. In contrast, the next-largest *R*^2^ for this relationship was 0.86 for dataset 3 and the average *R*^2^ for all datasets was 0.40 (see Table S2).

**Fig. 4.**
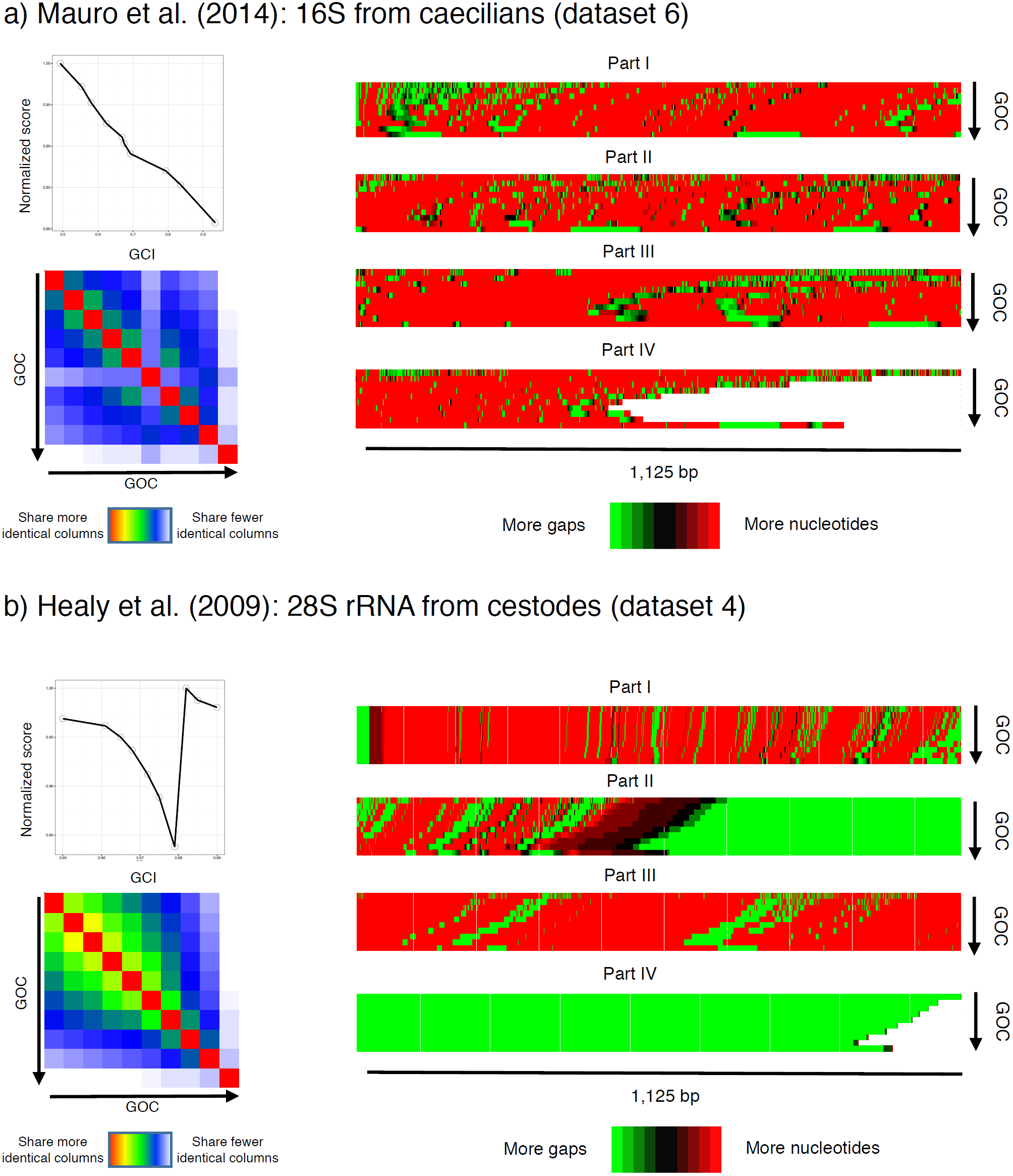
As an example of regular datasets, we show in a) the relationship of mean gap contiguity index (GCI) and the normalized likelihood scores (top left), the variation on the number of shared characters and the gap opening cost (GOC) (heatmap on the bottom left), and the distribution of gaps and nucleotides on all alignments (on the right) for datasets 6 (Mauro et al. 2014: 16S rRNA). In b) we show the same information for datasets 4 (Healy et al. 2009: 28S rRNA), which differs from all others on how indels affect the homology statements that follow them. The insertion of longer indels as the gap opening cost increases seems to result in increase in random homology statements as nucleotides are shifted to the right side of the alignment, leading to the unpredictability of mean GCI values. This also decreases similarity among characters in each alignment, and results in alignments that differs more in the character statements than on the distribution of gaps.

Although the average strength of the correlations between the aforementioned variables and the ML score was smaller than the correlation between gap opening cost and the ML score, we have no reason to assume the correlations are purely coincidental and instead propose that these variables partially account for the changes in the alignment matrix that lead to different ML scores. A special case seems to be when alignment is biased towards randomizing homology statements that follow long indels, as exemplified by dataset 4. In this case, the correlation of variables that explain the number and distribution of gaps in the alignment matrix with the ML score is weak, but we observed a strong correlation of the ML score and the TGI as a result of the number of gap↔nucleotide transformations on the tree.

### Fixed Nucleotide Homology and Alignment Length

When we fixed the nucleotide homologies and alignment lengths and varied the quantity and distribution of gaps by random nucleotide-to-gap replacements, we observed a strong, positive, linear relationship between the number of gaps (which in here vary in direct proportion to the probability replacing a nucleotide for an indels) and the ML score that was not dependent on the distribution of gaps in the alignment matrix (i.e., there was no correlation between the ML score and the any of the indices we defined, such as mean SGI; see Fig. 5).

**Fig. 5.**
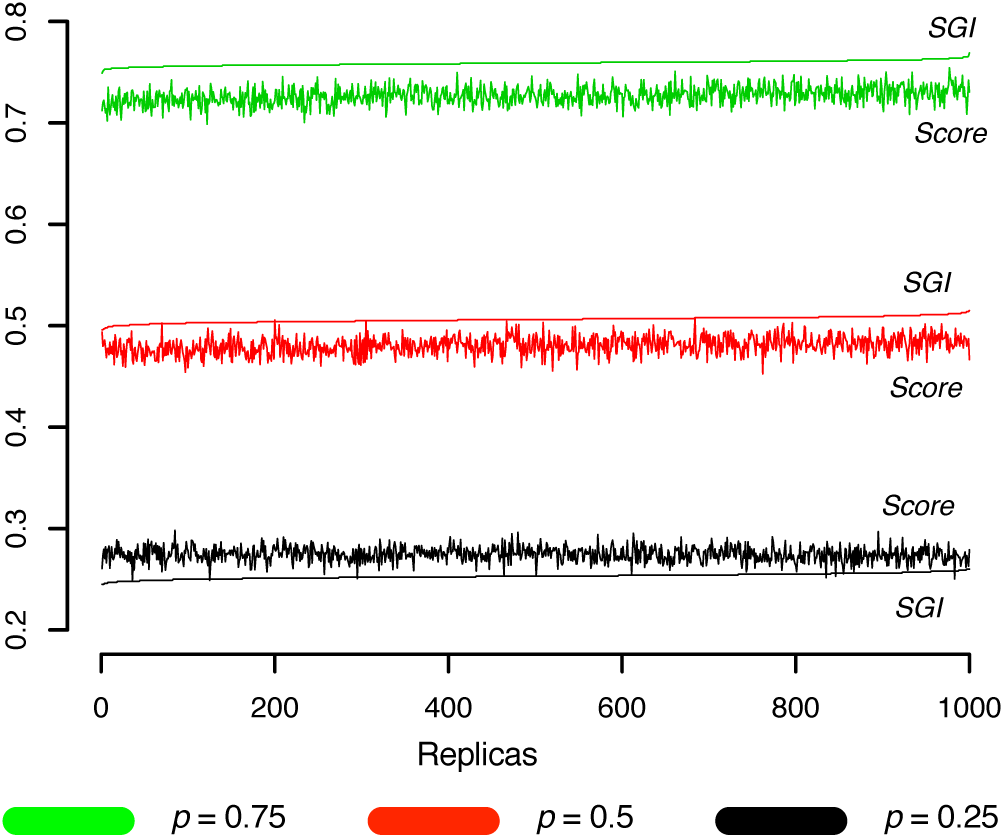
Variation of normalized likelihood score (LS) and shared gaps index (SGI) of dataset 8 across three rounds of simulations (1,000 independent replicas each). In each simulation round, nucleotides were substituted by indels with a fixed probability (black = 0.25, red = 0.5, and green = 0.75).

## Discussion

A long and growing list of theoretical and empirical studies has demonstrated the importance of treating gaps as indel events in phylogenetic analyses (e.g., 31–36). Nevertheless, the most common approach of practitioners is to code gaps as nucleotides of unknown identity (“N” or “?”). For example, our survey of this journal for the period 2013–2016 revealed that all published papers that performed a phylogenetic analysis of DNA sequences treated gaps as unknown nucleotides (studies that did not state how gaps were treated used methods that did not allow indel models). Given the predominance of this approach, it is imperative to understand the consequences of treating gaps as missing data in empirical studies.

Warnow et al. (37) demonstrated that ML phylogenetic methods of analysis are statistically inconsistent in certain conditions when gaps are treated as missing data, even when the true alignment is presented. More recently, Truszkowski and Goldman (38) demonstrated that “maximum likelihood phylogenetic inference is statistically consistent on multiple sequence alignments even in the case where gaps are treated as missing data, as long as substitution rates across edges are non-zero.” Truszkowski and Goldman (38) acknowledged that maximum likelihood is inconsistent in the scenario proposed by Warnow but argued that this scenario “is not representative of the vast majority of datasets where substitutions are common.” Neither of those studies addressed the behavior of empirical datasets analyzed using standard methods. This study is the first to systematically explore how the quantity and distribution of gaps treated as unknown nucleotides affect model selection, ML score, and topology in empirical phylogenetic analyses. Our general finding is that the effects depend on both the number of gaps and their effect on nucleotide↔nucleotide homologies. That is, all else being equal, as shown in our Monte Carlo simulations that randomly replaced nucleotides with gaps, increasing gaps results in higher ML scores and alignments approach trivial identity alignment (TIA; see 12). However, in practice, introducing more gaps during alignment also affects the homology relationship among nucleotides, resulting in less predictable outcomes.

On the basis of our results, we identify three general responses to variation in the number and distribution of gaps. The first response, exemplified by analyses of datasets 1–3 and 5–7, occurs when sequence length is similar among all terminals and variation in the number and distribution of gaps has little effect on nucleotide↔nucleotide homologies. In this scenario, ML scores are negatively correlated with gap opening cost, number of gaps, sequence length, and mean GCI, positively correlated with mean SGI, and unrelated to mean TGI. The second response is observed in analyses of dataset 4, which has the greatest variation in sequence length (Table 1). In this response, ML score is negatively correlated with gap opening cost and positively correlated with mean TGI, which suggests that the insertion of long indels in these sequences strongly affected nucleotide homology. Finally, the third response is drawn from our Monte Carlo simulations, whereby we introduced gaps into the alignment matrix without altering nucleotide homology. In this case, ML scores improve as gaps increasingly replaced nucleotides, showing that, all else being equal, ML score increases with the amount of missing data (cf. 12).

In both responses 1 and 2, the uniformity of the models of nucleotide evolution selected for the different alignments was unexpected. Given that alignments approaching TIA are simple datasets that need few transformations due to maximization of character columns that include only one nucleotide class (i.e., identical nucleotides and gaps treated as nucleotides of unknown identity), we expected that gappier alignments would require less complex models. Our interpretation of the lack of variation in model selection is that the alignments did not sufficiently approximate TIA to reduce the complexity of the models needed to explain the data. This explanation is supported by the fact that the most parameter-rich model was selected as optimal (i.e., GTR+I+G) for most alignments (75%).

We caution that our findings are agnostic with regards to the optimal gap opening and extension costs for empirical analyses. That is, although the effects of variation in gap costs on model selection, tree topology, and ML score can be predicted, none of these response variables provides a defensible optimality criterion for selecting alignments or alignment parameters in standard ML analysis. The program SATé (39, 40) does employ ML score as an optimality criterion to choose among alignments obtained from MAFFT, but Denton and Wheeler (12) showed that the gaps-as-missing assumption results in TIA being optimal if alignments are evaluated on the basis of the ML score. In practice, it is highly unlikely for trivial alignments to be chosen as optimal in empirical studies because SATé searches using alignments obtained from MAFFT, which does not use ML as its optimality criterion and does not treat gaps as absence of evidence. Nevertheless, this does not absolve SATé of Denton and Wheeler’s (12) fundamental criticism, as its apparent immunity is due to its incomplete analysis of alignment space and inconsistent application of the optimality criterion. That is, given the specified optimality criterion, an adequately thorough analysis must select TIA as optimal, and it is only by employing different criteria for alignment and tree assessment that SATé avoids TIA. As Denton and Wheeler (12) demonstrated, the problem is eliminated if gaps are attributed a cost in both the alignment and tree searching stages of analysis, which has the further advantage of allowing the same optimality criterion to be applied through generalized tree-alignment (41), as envisioned by (42) and implemented in programs like POY (43) and BEAST (44).

## Supplementary Material

Supplementary material, including original datasets, home-made scripts, and Tables S1-2, is available at zenodo.org, doi: 10.5281/zenodo.3363990.

## Acknowledgements

Funding was provided by the Fundação de Amparo à Pesquisa do Estado de São Paulo (Grant Numbers: 2012/10000-5, 2013/09598-8, 2015/18654-2, 2018/15425-0) and Conselho Nacional de Desenvolvimento Científico e Tecnológico (Grant Number: 306823/2017-9). We thank the organizers of the 35th Annual Meeting of the Willi Hennig Society and XII Reunión Argentina de Cladística y Biogeografia (Museo Argentino de Ciencias Naturales “Bernardino Rivadavia”, Buenos Aires, Argentina, October 5-8, 2016) where this work was first presented. The development of the ideas presented in this study benefited from discussions with Darrel Frost, John Denton, Jose Padial, Pedro Peloso, and Ward Wheeler. We also thank Pedro Ivo Simões for his criticisms of previous versions of the manuscript.

